# ColabFold - Making protein folding accessible to all

**DOI:** 10.1101/2021.08.15.456425

**Authors:** Milot Mirdita, Konstantin Schütze, Yoshitaka Moriwaki, Lim Heo, Sergey Ovchinnikov, Martin Steinegger

## Abstract

ColabFold offers accelerated protein structure and complex predictions by combining the fast homology search of MMseqs2 with AlphaFold2 or RoseTTAFold. ColabFold’s 40 - 60× faster search and optimized model use allows predicting close to a thousand structures per day on a server with one GPU. Coupled with Google Colaboratory, ColabFold becomes a free and accessible platform for protein folding. ColabFold is open-source software available at github.com/sokrypton/ColabFold. Its novel environmental databases are available at colabfold.mmseqs.com

---

Predicting the three-dimensional structure of a protein from its sequence alone remains an unsolved problem. However, by exploiting the information in multiple sequence alignments (MSAs) of related proteins as raw input features for end-to-end training, AlphaFold2 [1] was able to predict the 3D atomic coordinates of folded protein structures at a median GDT-TS of 92.4% in the latest CASP14 [2] competition. The accuracy of many of the predicted structures was within the error margin of experimental structure determination methods. Many ideas of AlphaFold2 were independently reproduced and implemented in RoseTTAFold [3]. Additionally to single chain predictions, RoseTTAFold was shown to model protein complexes. Evans *et al*. [4] released AlphaFold-multimer, a refined version of AlphaFold2 for complex prediction. Thus, two highly accurate open-source prediction methods are now publicly available.

In order to leverage the power of these methods researchers require powerful compute-capabilities. First, to build diverse MSAs, large collections of protein sequences from public reference [5] and environmental [1, 6] databases are searched using the most sensitive homology detection methods HMMer [7] and HHblits [8]. These environmental databases contain billions of proteins extracted from metagenomic and -transcriptomic experiments, which often complement databases dominated by isolate genomes. Due to their large size searches can take up to hours for a single protein, while requiring over two terabyte of storage space alone. Second, to execute the deep neural networks GPUs with a large amount of GPU RAM are required even for relatively common protein sizes of ~1000 residues. Though, for these the MSA generation dominates the overall run-time.

To enable researchers without these resources to use AlphaFold2, independent solutions based on Google Colaboratory were developed. Colaboratory is a proprietary version of Jupyter Notebook hosted by Google. It is accessible for free to logged-in users and includes access to powerful GPUs. Tunyasuvunakool *et al*. [9] developed an AlphaFold2 Jupyter Notebook for Google Colaboratory (referred to as AlphaFold-Colab), where the input MSA is built by searching with HMMer against a clustered UniProt and an eight-fold reduced environmental databases. Resulting in less accurate predictions, while still requiring long search times.

Here, we present ColabFold, a fast and easy to use software for protein structure and homo- and heteromer complex prediction, for use as a Jupyter Notebook inside Google Colaboratory, on researchers’ local computers as a notebook or through a command line interface. ColabFold speed-ups the prediction by replacing AlphaFold2’s homology search with a 40-60 times faster MMseqs2 [10, 11] search. It additionally implements speed-ups for batch predictions of structures by avoiding recompilation and adding early stop criteria. ColabFold’s batch mode with early stopping can compute the proteome of *Methanocaldococcus jannaschii* in 48 h on a consumer GPU a ~90 times speedup over AlphaFold2. We show that Colab-Fold outperforms AlphaFold-Colab and matches AlphaFold2 on CASP14 targets and also matches AlphaFold-multimer on the ClusPro [4, 12] dataset in prediction quality.

ColabFold (**Fig. 1**) consists of three parts: (1) An MMseqs2 based homology search server to build diverse MSAs and to find templates. The server efficiently aligns input sequence(s) against the UniRef100, the PDB70 and an environmental sequence set. (2) A Python library that communicates with the MMseqs2 search server, prepares the input features for (single or complex) structure inference, and visualizes of results. This library also implements a command line interface. (3) Jupyter notebooks for basic, advanced and batch use (Methods “ColabFold notebooks”) using the Python library.

**FIG. 1.**
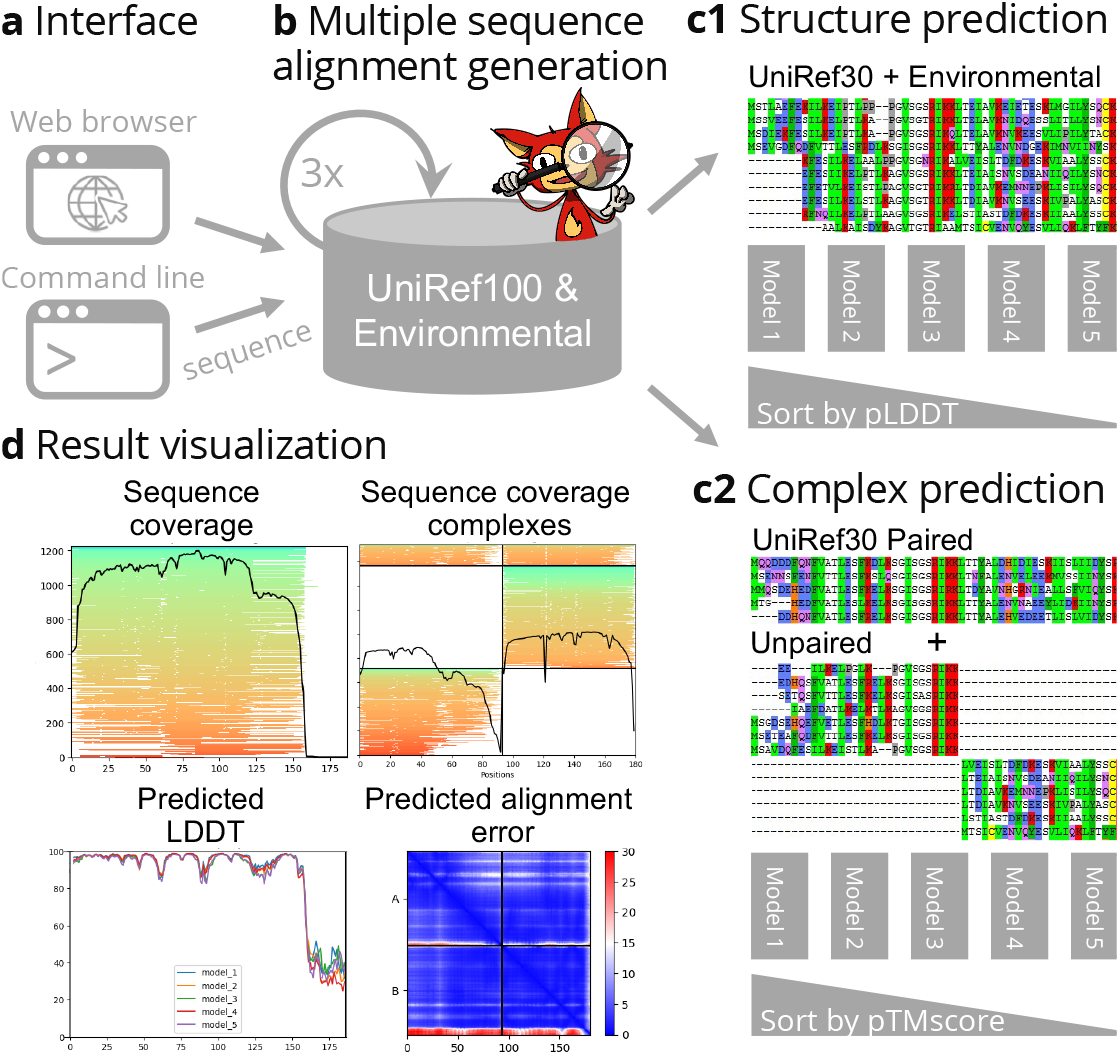
(**a**) ColabFold has a web and a command line interface, that (**b**) send FASTA input sequence(s) to a MMseqs2 server searching two databases UniRef100 and a database of environmental sequences with three profile-search iterations each. The second database is searched using a sequence-profile generated from the UniRef100 search as input. The server generates two MSAs in A3M format containing all detected sequences. (**c1**) For single structure predictions we filter both A3Ms using a diversity aware filter and return this to be provided as the MSA input feature to the AlphaFold2 models. (**c2**) For complex prediction we pair the top hits within the same species to resolve the inter-complex contacts and additionally add two unpaired MSAs (same to **c1**) to guide the structure prediction. (**d**) To help researchers judge the prediction quality we visualize MSA depth and diversity and show the AlphaFold2 confidence measures (pLDDT and PAE).

In ColabFold we replace the sensitive search methods HMMer and HHblits by MMseqs2. We optimized the MSA generation by MMseqs2 to have the following three properties: (1) MSA generation should be fast. (2) The MSA has to capture diversity well and (3) it has to be small enough to run on computers with limited RAM. Reducing the memory requirement is especially helpful in Google Colaboratory where the provided system is selected from a pool with widely differing capabilities. While (1) is achieved through the fast MMseqs2 prefilter for (2 and 3) we developed a search workflow to maximize sensitivity (Methods “MSA generation”) and a new filter that samples the sequence space evenly (Methods “New diversity aware filter” and **Supplementary Fig. 1**). Prediction quality highly depends on the input MSA. However, often an MSA with only a few (~30) sufficiently diverse sequences is enough to produce high quality predictions (see Jumper et al., **Fig. 5a**).

Additionally, we combined the BFD and MGnify databases that are used in AlphaFold2 by HHblits and HMMer respectively into a combined redundancy reduced version we refer to as BFD/MGnify (Methods “Reducing size of BFD/MGnify”). The environmental search database presented an opportunity to improve structure predictions of non-bacterial sequences, as e.g., eukaryotic protein diversity is not well represented in the BFD and MGnify databases. Limitations in assembly and gene calling due to complex intron/exon structures result in under representation in reference databases. We therefore extended the BFD/MGnify with additional metagenomic protein catalogues containing eukaryotic proteins [13, 14, 15], phage catalogues [16, 17] and an updated version of MetaClust [18]. We refer to this database as ColabFoldDB (Methods “Colab-FoldDB”). In **Supplementary Fig. 2** we show that the ColabFoldDB in comparison to the BFD/MGnify produces more diverse MSAs for PFAM [19] domains with < 30 members.

To compare the accuracy of predicted structures we compared AlphaFold2 (default settings with templates), AlphaFold-Colab (no templates), ColabFold-RoseTTAFold-BFD/MGnify, ColabFold-AlphaFold2-BFD/MGnify and ColabFold-AlphaFold2-ColabFoldDB on TM-scores for all targets from the CASP14 competition (**Fig. 2a**). All three ColabFold modes were executed without templates. We show the targets split by free modeling (FM) on the left and the remaining ones on the right, since we used the FM-targets for optimization of search workflow parameters.

**FIG. 2.**
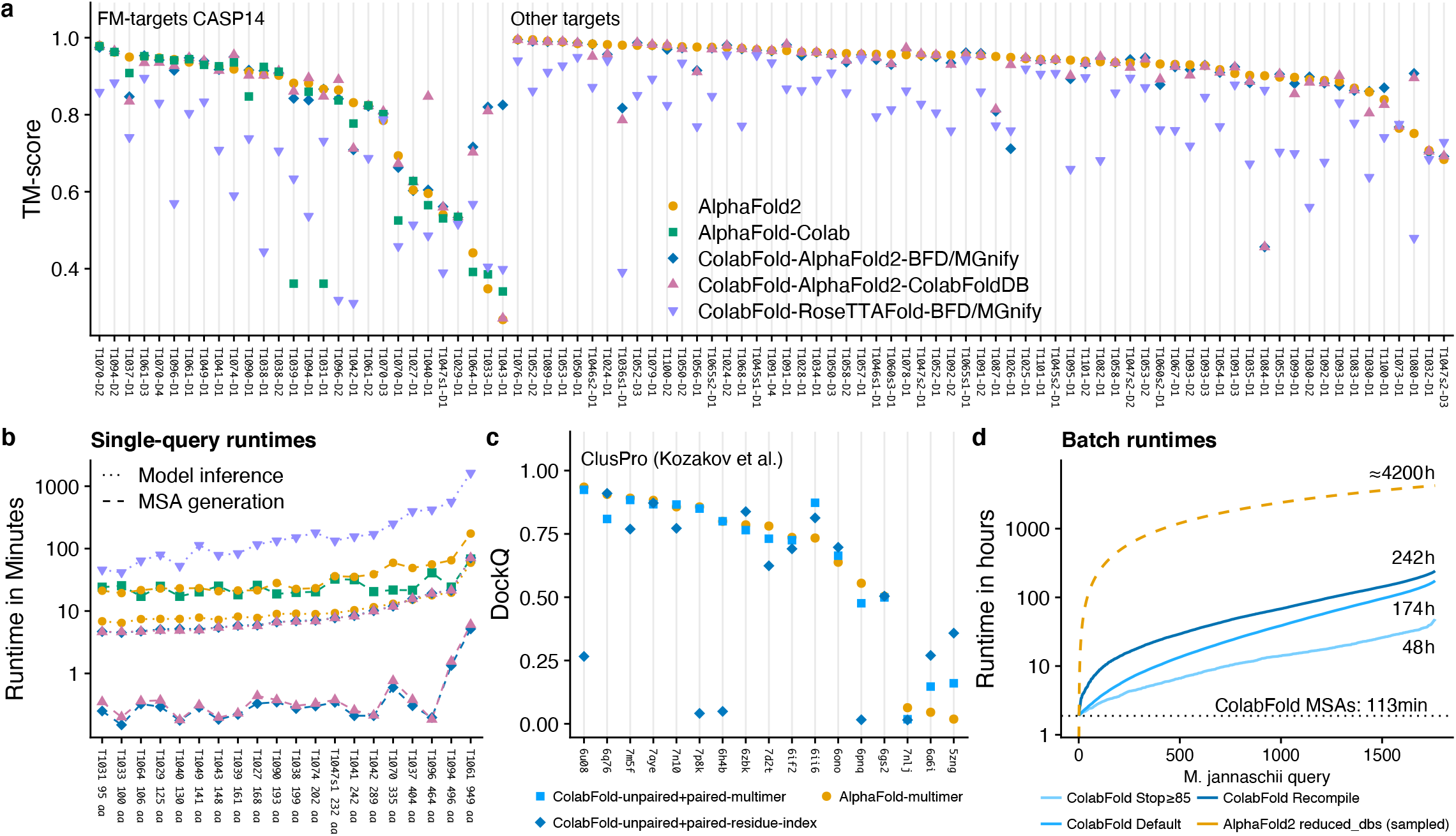
(**a**) Structure prediction comparison of AlphaFold2 (yellow), AlphaFold-Colab (green) and ColabFold-AlphaFold2 with BFD/MGnify (blue) and with the ColabFoldDB (magenta), and ColabFold-RoseTTAFold with BFD/MGnify (purple) using predictions of 91 domains of 65 CASP14 targets. The 28 domains from the 20 free-modeling (FM) targets are shown first. FM targets were used to optimize MMseqs2 search parameters. Each target was evaluated for each individual domain (in total 91 domains). (**b**) MSA generation and model inference times for each CASP14 FM target sorted by protein length (same colors as before). Blue shows MSA runtimes for ColabFold-AlphaFold2-BFD/MGnify and ColabFold-RoseTTAFold-BFD/MGnify. (**c**) Comparison of ColabFold complex predictions in residue-index-(dark blue) and AlphaFold-multimer (light blue) mode, and to AlphaFold-multimer (yellow). (**d**) Runtime of colab-fold_batch proteome prediction at three optimization levels: (dark blue) Always recompile, (blue) default, (light blue) stop model/recycle evaluation after first prediction with a pLDDT of *≥*85. Extrapolated line based on 50 AlphaFold2 predictions shown in yellow.

The mean TM-scores for the FM targets are 0.826, 0.818, 0.79, 0.744 and 0.62 for ColabFold-AlphaFold2-BFD/MGnify, ColabFold-AlphaFold2-ColabFoldDB, AlphaFold2, AlphaFold-Colab and ColabFold-RoseTTAFold-BFD/MGnify respectively. Over all CASP14 targets the TM-scores are 0.887, 0.886, 0.888 and 0.754 for the respective methods, excluding AlphaFold-Colab as it cannot be used stand-alone.

ColabFold could not predict T1084 well as MMseqs2 suppresses all databases hits as false positives due to its amino acid composition filter and masking procedure. If these filters are deactivated T1084 can be predicted with an TM-score of 0.872 (**Supplementary Fig. 3**). **Supplementary Table 1** contains a list of further targets where ColabFold differed significantly from AlphaFold2.

ColabFold is on average 5x faster for single predictions than AlphaFold2 and AlphaFold-Colab, when taking both MSA generation (**Fig. 2b**) and model inference into account.

AlphaFold2 was initially released without capabilities to model complexes. However, we found that by combining two sequences with a glycine linker [20] it could often successfully model complexes. Shortly afterwards, Baek [21] found that incrementing the model-internal residue index - the method that was used in RoseTTAFold - could also be used in AlphaFold2.

For high quality predictions it was shown that sequences should be provided in paired-form to AlphaFold2 [22]. We implemented a similar pairing procedure (Methods “MSA pairing for complex prediction”) and show the complex prediction capabilities of ColabFold in **Fig. 2c**. ColabFold achieves the highest accuracy in complex prediction on the ClusPro [4, 12] dataset with the AlphaFold-multimer model, however, some targets performed better using the residue-index mode.

**Fig. 3** shows two examples of ColabFold’s complex prediction capabilities: (**a**) shows a homo-six-mer and (**b**) shows a D-methionine transport system composed of three different proteins. For single structure prediction AlphaFold2 provides a pLDDT measure to indicate the prediction quality. A high pLDDT does not necessarily indicate a correct complex prediction, though the inter-complex predicted alignment error (PAE) helps to rank complexes. We visualize plots of PAE and complex conformation to help users judge the prediction quality of a complex. An example for heteromer complex prediction is shown in **Supplementary Fig. 4** with its PAE plot. Furthermore, ColabFold complexes were successfully used to aid the cryo-EM structure determination of the 120 MDa human nucleopore complex [23].

**FIG. 3.**
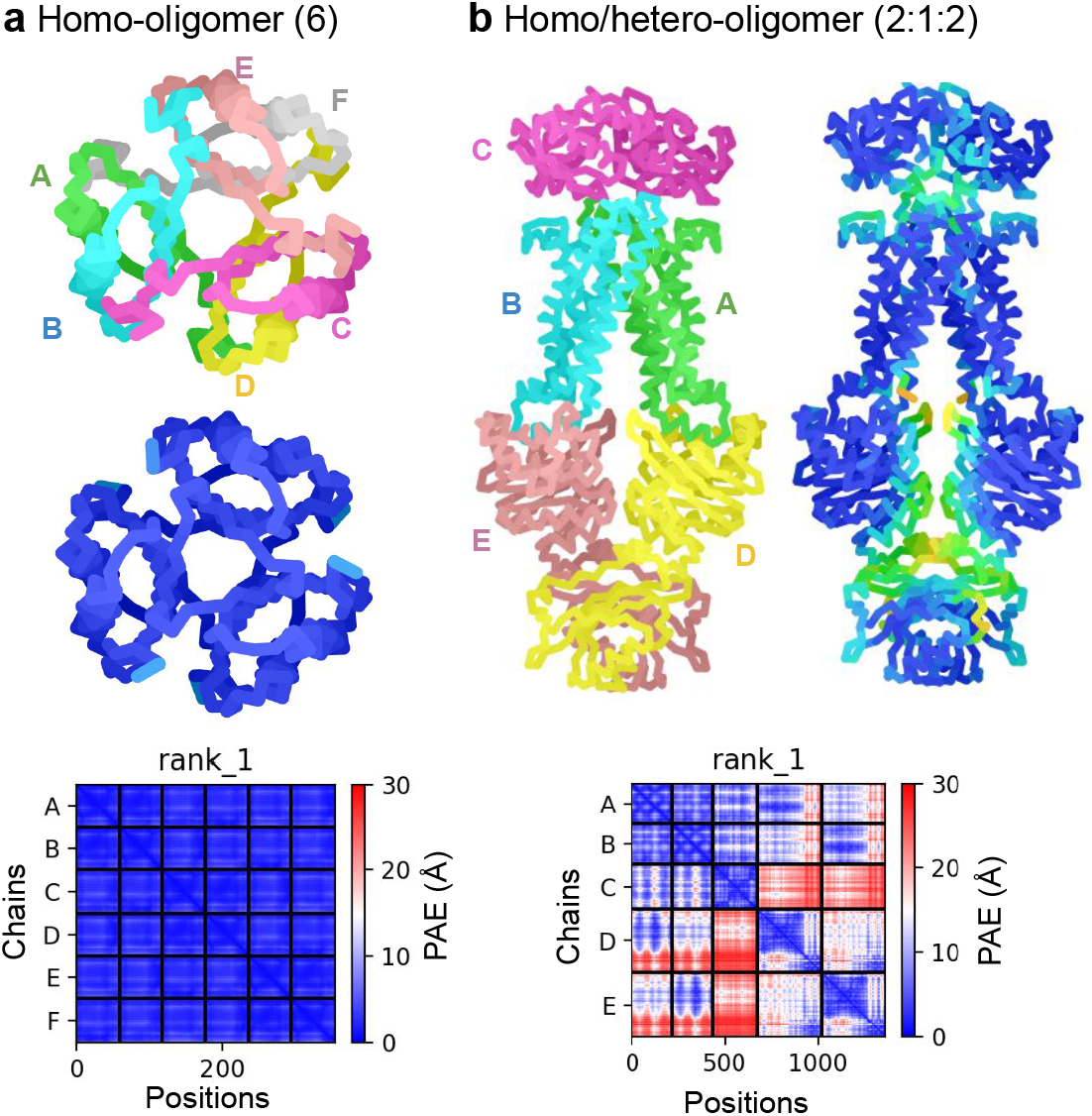
Anecdotal examples showcasing the capabilities of advanced ColabFold features. (**a**) Setting the homo-oligomer setting to 6, allows modeling of the homo-6-mer structure of 4-Oxalocrotonate Tautomerase. Colored by chain (top), pLDDT (predicted Local Distance Difference Test, bottom). The inter PAE (Predicted Aligned Error) between chains is very low indicating a confident prediction. (**b**) Providing three different proteins with 2:1:2 homo-oligomer setting allows modeling a hetero-complex with mismatching symmetries of the D-methionine transport system.

In ColabFold we expose many internal parameters of AlphaFold2 to aid users to model difficult targets, such as the recycle count (default 3). It controls the number of times the prediction is repeatedly fed through the model. For difficult targets as well as for designed proteins without known homologs additional recycling iterations can result in a high quality prediction (**Supplementary Fig. 5**). Rerunning the CASP14 benchmark using 12 recycles resulted in an improvement of average TM-score from 0.887 to 0.898 (**Supplementary Fig. 6**). The largest improvement was in targets with little MSA information.

To meet the demand for high throughput structure prediction we introduced several features in ColabFold. (1) MSA generation can be executed in batch-mode independently from model batch-inference. (2) We compile only one of the five AlphaFold2 models and reuse weights. (3) We provide a batch execution mode, that avoids recompilation for sequences of similar length. (4) We implement early stop criteria, to avoid running additional recycles or models if a sufficiently accurate structure was already found. (5) We developed the command line tool `colabfold_batch` to predict structures on local machines. All together, we show that the proteome of 1762 proteins shorter than 1000 aa of *M. jannaschii* can be predicted in 48 h with early stopping at pLDDT of ≥85 on one Nvidia Titan RTX (**Fig. 2d**), while sacrificing little-or-no prediction accuracy (Methods “Proteome Benchmark”). The average pLDDTs of AlphaFold2 and ColabFold Stop ≥ 85 were 89.75 and 88.78 in a subsampled set of 50 proteins.

ColabFold builds beyond the initial offerings of Alphafold2 by improving its sequence search, providing tools for modeling homo- and heteromer complexes, exposing advanced functionality, expanding the environmental databases and enabling large-scale batch prediction of protein structures – at a ~90 times speedup over AlphaFold2.

## Acknowledgements

We thank Johannes Söding for providing computational resources. Richard Evans, John Jumper, and Tim Green for answering questions regarding AF2. Minkyung Baek for the complex residue trick. Do-Yoon Kim for creating the ColabFold logo. Enzo Guerrero-Araya and Jakub Kaczmarzyk for providing bug fixes. Alon Markovich and Julia Varga for notifying us about MSA quality issues. Harriet Alexander for providing the TOPAZ proteins as a single file to download. We thank all users for using ColabFold and reporting issues.

This work used the Scientific Compute Cluster at GWDG, the joint data center of Max Planck Society for the Advancement of Science (MPG) and University of Göttingen. Milot Mirdita acknowledges the BMBF CompLifeSci project horizontal4meta. Martin Steinegger acknowledges support from the National Research Foundation of Korea grant [2019R1A6A1A10073437, 2020M3A9G7103933, 2021R1C1C102065, 2021M3A9I4021220]; New Faculty Startup Fund and the Creative-Pioneering Researchers Program through Seoul National University. Yoshitaka Moriwaki acknowledges support from Platform Project for Supporting Drug Discovery and Life Science Research (Basis for Supporting Innovative Drug Discovery and Life Science Research (BINDS)) from AMED under Grant Number JP21am0101107. For this project, Sergey Ovchinnikov was supported by the National Science Foundation under Grant No. MCB2032259. Any opinions, findings, and conclusions or recommendations expressed in this material are those of the author(s) and do not necessarily reflect the views of the 220 National Science Foundation.

## AUTHOR CONTRIBUTION

M.M., K.S., S.O. and M.S. performed research and program-222 ming, M.M., S.O. and M.S. jointly designed the research and 223 wrote the manuscript. Y.M. provided the initial methodology 224 for hetero-complex modeling and created an installer for use 225 on local servers. L.H. provided initial benchmarking.

## MATERIALS AND METHODS

### Executing ColabFold

ColabFold is available as a set of Jupyter notebooks, to use on Google Colaboratory or users’ local machines, as well as an easily installable command line application.

#### ColabFold notebooks

ColabFold has four main Jupyter notebooks [24]: `AlphaFold2_mmseqs2` for basic use that supports protein structure prediction using (1) MSAs generated by MMseqs2, (2) custom MSA upload, (3) using template information, (4) relaxing the predicted structures using amber force fields [25], and (5) complex prediction. `AlphaFold2_advanced` for advanced users additionally supports (6) MSA generation using HMMer (same as AlphaFold-Colab), (7) the sampling of diverse structures by iterating through a series of random seeds (`num_samples`), and (8) control of AlphaFold2 model internals, such as changing the number of recycles (`max_recycle`), number of ensembles (`num_ensemble`), and enabling the stochastic part of the models via the (`is_training`) option. The latter enables dropout during inference, allowing the user to sample solutions from the uncertainty of the model [26] or the ambiguity of co-evolution constraints derived from the input MSA. `AlphaFold2_batch` for batch prediction of multiple sequences or MSAs. The batch notebook saves time by avoiding recompilation of the AlphaFold2 models (“Avoid recompiling during batch computation”) for each individual input sequence. `RoseTTAFold` for basic use of RoseTTAFold that supports protein structure prediction using (1) MSAs generated by MM-seqs2, (2) custom MSAs and (4) sidechain prediction using SCWRL4 [27]. The RoseTTAFold notebook also has an option use a slower but more accurate PyRosetta [28] folding protocol for structure prediction, using constraints predicted by RoseTTAFold’s neural network.

#### ColabFold command line interface

We initially focused on making ColabFold as widely available as possible through our Notebooks running in Google Colaboratory. To meet the demand for a version that runs on local users’ machines, we released “LocalColabFold”. LocalColabFold can take command line arguments to specify an input FASTA file, an output directory, and various options to tweak structure predictions. LocalColabFold runs on wide range of operating systems, such as Windows 10 or later (using Windows Subsystem for Linux 2), macOS, and Linux. The structure inference and energy minimization are accelerated if a CUDA 11.1 or later compatible GPU is present. LocalColabFold is available as free open-source software at github.com/YoshitakaMo/localcolabfold.

Recognizing the limitations of Google Colaboratory, we provide the `colabfold_batch` command line tool through the `colabfold` python package. This allows computing of tasks too large for Google Colab on users’ own computer, e.g. predicting an entire proteome (Methods “Proteome benchmark”). It can be installed with `pip install colabfold, followed by pip install -U “jax[cuda]” -f` https://storage.googleapis.com/jax-releases/jax_releases.html. It can be used as `colabfold_batch input_file_or_directory output_directory`, supporting FASTA, A3M and CSV files as input.

### Replacing MSA generation in AlphaFold2/RoseTTAFold with MMseqs2

Generating multiple sequence alignments for AlphaFold2 and RoseTTAFold is a time-consuming task. To improve their runtime, while maintaining a high prediction accuracy, we implemented optimized workflows using MMseqs2.

#### MSA generation by MMseqs2

ColabFold sends the query sequence to a MMseqs2 server [11]. It searches the sequence(s) with three iterations against the consensus sequences of the UniRef30, a clustered version of the UniRef100 [29]. We accept hits with an E-value of lower than 0.1. For each hit, we realign its respective UniRef100 cluster member using the profile generated by the last iterative search, filter them (Methods “New diversity aware filter”) and add these to the MSA. This expanding search results in a speed up of ~10x as only 29.3 million cluster consensus sequence are searched instead of all 277.5 million UniRef100 sequences. Additionally, it has the advantages to be more sensitive since the cluster consensus sequences are used. We use the UniRef30 sequence-profile to perform an iterative search against the BFD/MGnify or ColabFoldDB using the same parameters, filters and expansion strategy.

#### New diversity aware filter

To limit the number of hits in the final MSA we use the HHblits diversity filtering algorithm [8] implemented in MMseqs2 in multiple stages: (1) During UniRef cluster expansion, we filter each individual UniRef30 cluster before adding the cluster members to the MSA, such that no cluster-pair has a higher maximum sequence identity than 95% (`--max-seq-id 0.95.` (2) After realignment enable only the `-- qsc 0.8` threshold and disable all other thresholds (`--qid 0 --diff 0 --max-seq-id 1.0`). Additionally, the qsc filtering is only used if least 100 hits were found (`--filter-min-enable 100`). (3) During MSA construction we filter again with the following parameters: `--filter-min-enable 1000 --diff 3000 --qid 0.0,0.2,0.4,0.6,0.8,1.0 --qsc 0 --max-seq-id 0.95.` Here, we extended the HHblits filtering algorithm to filter within a given sequence identity bucket, such that it cannot eliminate redundancy across filter buckets. Our filter keeps the 3000 most diverse sequences in the identity buckets ]0.0-0.2], ]0.2-0.4], ]0.4-0.6], ]0.6-0.8] and ]0.8-1.0]. In buckets containing less than 1000 hits we disable the filtering.

#### New MMseqs2 pre-computed index to support expanding cluster members

MMseqs2 was initially built to perform fast many-against-many sequence searches. Mirdita *et al*. improved it to also support fast single-against-many searches. This type of search requires the database to be index and stored in memory `mmseqs createindex` indexes the sequences and stores all time-consuming-to-compute data structures used for MMseqs2 searches to disk. We load the index into the operating systems cache using *vmtouch* (github.com/hoytech/vmtouch) to allow calls to the different MMseqs2 modules to become near-overhead free. We extended the index to store, in addition to the already present cluster consensus sequences, all member sequences and the pairwise alignments of the cluster representatives to the cluster members. With these resident in cache, we eliminate the overhead of the remaining module calls.

#### ColabFold databases

AlphaFold2 requires over 2 terabyte of storage space for its databases, which is a significant hurdle for many researchers. We optimized its databases and additionally created another large environmental sequence database.

#### Reducing size of BFD/MGnify

To keep all required sequences and data structures in memory we needed to reduce the size of the environmental databases BFD and MGnify, as both databases together would have required ~517 GB RAM for headers and sequences alone.

BFD is a clustered protein database consisting of ~2.2 billion proteins organized in 64 million clusters. MGnify (2019_05) contains ~300 million environmental proteins. We merged both databases by searching the MGnify sequences against the BFD cluster representative sequences using MMseqs2. Each MGnify sequence with a sequence identity of >30% and a local alignment that covers at least 90% of its length is assigned to the respective BFD cluster. All unassigned sequences are clustered at >30% sequence identity and 90% coverage (`--min-seq-id 0.3 -c 0.3 --cov-mode 1 -s 3`) and merged with the BFD clusters, resulting in 182 million clusters. In order to reduce the size of the database we filtered each cluster keeping only the 10 most diverse sequences using (`mmseqs filterresult --diff 10`). This reduced the total number of sequences from 2.5 billion to 513 million, thus requiring only 84 GB RAM for headers and sequences.

### ColabFoldDB

We built ColabFoldDB by expanding the BFD/MGnify with metagenomic sequences from various environments. To update the database, we searched the proteins from the SMAG (eukaryotes) [14], MetaEuk (eukaryotes) [13], TOPAZ (eukaryotes) [15], MGV (DNA viruses) [16], GPD (bacteriophages) [17] and updated version of MetaClust [18] against the BFD/MGnify centriods using MMseqs2 and assigned each sequence to the respective cluster if they have a 30% sequence identity at a 90% sequence overlap (`-c 0.9 --cov-mode 1 --min-seq-id 0.3`). All remaining sequences were clustered using MMseqs2 `cluster -c 0.9 --cov-mode 1 --min-seq-id 0.3` and appended to the database. We remove redundancy per cluster by keeping the most 10 diverse sequences using (`mmseqs filterresult --diff 10`). The final database consists of 209,335,865 million representative sequences and 738,695,580 members. See “Data availability” for input files. We provide the MMseqs2 search workflow used in the server (“MSA generation by MMseqs2”) as a standalone script `colabfold_search.sh`.

#### Template information

AlphaFold2 searches with HHsearch through a clustered version of the PDB (PDB70 [8]) to find the 20 top ranked templates. In order to save time, we use MMseqs2 [10] to search against the PDB70 cluster representatives as a prefiltering step to find candidate templates. This search is also done as part of the MMseqs2 API call on our server. Only the top 20 target templates according to E-value are then aligned by HHsearch. The accepted templates are given to AlphaFold2 as input features. This alignment step is done in the ColabFold client and therefore requires the subset of the PDB70 containing the respective HMMs. The PDB70 subset and the PDB mmCIF files are fetched from our server. For benchmarking, no templates are given to ColabFold.

### Modeling protein complexes with ColabFold

ColabFold offers protein complex folding through the specialized AlphaFold-multimer model and through residue-index manipulation [3]. Here, we show the steps we took for ColabFold to produce accurate protein complex predictions.

#### Modeling of protein-protein complexes

We implemented two protein complex prediction modes in ColabFold. One based on AlphaFold-multimer [4] and one based on the residue index manipulation of the original AlphaFold2 model. Baek *et al*. [3] show that RoseTTAFold is able to model complexes, despite being trained only on single chains. This is done by providing a paired alignment and modifying the residue index. The residue index is used as an input to the models to compute positional embeddings. In AlphaFold2, we find the same to be true, although surprisingly the paired alignment is often not needed (**Fig. 2c**). AlphaFold2 uses relative positional encoding with a cap at *|i – j|* ≥ 32. Meaning, any pair of residues separated by >32 or more are given the same relative positional encoding. By offsetting the residue index between two proteins to be > 32, AlphaFold2 treats them as separate poly-peptide chains. ColabFold integrates this for modeling complexes.

For homo-oligomeric complexes (**Fig. 3a**), the MSA is copied multiple times for each component. Interestingly, it was found that providing a separate MSA copy (padding by gap characters to extend to other copies) to work significantly better than concatenating left-to-right.

For hetero-oligomeric complexes (**Fig. 3b**), a separate MSA is generated for each component. The MSA is paired according to the chosen `pair_mode` (“MSA pairing for complex predic tion”). Since pLDDT is only useful for assessing local structure confidence, we use the fine-tuned model parameters to return the PAE for each prediction. As illustrated in **Supplementary Fig. 4**, the inter-PAE (predicted aligned error), the predicted TM-score or interface TM-score (both derived from PAE) can be used to rank and assess the confidence of the predicted protein-protein interaction.

#### MSA pairing for complex prediction

A paired MSA helps AlphaFold2 to predict complexes more accurately only if orthologous genes are paired with each other. We followed a similar strategy as Bryant *et al*. [22] to pair sequences according to their taxonomic identifier. For the pairing we search each distinct sequence of a complex against the UniRef100 using the same procedure as described in “MSA generation”. We return only hits that cover all complex proteins within one species and pair only the best hit (smallest e-value) with an alignment that covers the query to at least 50%. The pairing is implemented in the new MMseqs2 module `pairaln`.

For prokaryotic protein prediction, we additionally implemented the protocol described in [3] to pair sequences based on their distances in the genome as predicted from the UniProt accession numbers.

#### Taxonomic labels for MSA pairing

To pair MSAs for complex prediction, we retrieve for each found UniRef100 member sequence the taxonomic identifier from the NCBI taxonomy [30]. The taxonomic labels are extracted from the lowest common ancestor field (“common taxon ID”) of each UniRef100 sequence from the `uniref100.xml` (2021_03) file.

### Speeding up AlphaFold2’s model evaluation

Our efforts in speeding up AlphaFold2’s MSA generation yielded large improvements in its runtime. However, we discovered multiple opportunities within AlphaFold2 to speed up its model inference, without sacrificing (or only sacrificing very little) of its accuracy.

### Avoid recompiling AlphaFold2 models

The AlphaFold2 models are compiled using JAX [31] to optimize the model for specific MSA or template input sizes. When no templates are provided, we compile once and, during inference, replace the weights from the other models, using the configuration of model 5. This saves 7 minutes of compile time. When templates are enabled, model 1 is compiled and weights from model 2 are used, model 3 is compiled and weights from models 4 and 5 are used. This saves 5 minutes of compile time. If the user changes the sequence or settings, without changing the length or number of sequences in the MSA, the compiled models are reused without triggering recompilation.

#### Avoid recompiling during batch computation

In order to avoid AlphaFold2 model recompilation for every protein AlphaFold2 provides a function to add padding to the input MSA and templates called `make_fixed_size`. However, this is not exposed in AlphaFold2. We used the function in our batch notebook as well as in our command line tool *colabfold_batch*, in order to maximize GPU utilization and minimize the need of model recompilation. We sort the input queries by sequence length and process them in ascending order. We pad the input features by 10% (by default). All sequences that lie within the query length and an additional 10% margin do not require to be recompiled, resulting in a large speed up for short proteins.

#### Recycle count

AlphaFold2 improves the predicted protein structure by recycling (by default) 3 times, meaning the prediction is fed multiple times through the model. We exposed the recycle count as a customizable parameter as additional recycles can often improve a model (**Supplementary Fig. 6**) at the cost of a longer runtime. We also implemented an option to specify a tolerance threshold to stop early. For some designed proteins without known homologous sequences, this helped to fold the final protein (**Supplementary Fig. 5**).

#### Speed-up of predictions through early stop

AlphaFold2 computes five models through multiple recycles. We noted that for prediction of high certainty (*>* 85 pLDDT), all five models would often produce structures of very similar confidence, for some even without or with less than 3 of recycles. In order to speed up the computation we added a parameter to define an early stop criterion that halts additional model inferences and stops recycling if a given pLDDT or (interface) pTMscore threshold is reached.

#### Exposing advanced features

In our investigation of AlphaFold2’s internals, we realized that we could expose many knobs that might be usefully to researchers trying to explore AlphaFold2’s full potential.

#### Sampling of diverse structures

To reduce memory requirements, only a subset of the MSA is used as input to the model. Alphafold2, depending on model configuration, subsamples the MSA to a maximum of 512 cluster centers and 1024 “extra” sequences. Changing the random seed can result in different cluster centers and thus different structure predictions. ColabFold provides an option to iterate through a series of random seeds, resulting in structure diversity. Further structure diversity can be generated by using the original or fine-tuned (`use_ptm`) model parameters and/or enabling (`is_training`) to activate the stochastic (dropout) part of model. Enabling the latter, can be used to sample an ensemble of models for the uncertain parts of the structure prediction.

#### Custom MSAs

ColabFold allows researchers to upload their own MSAs. Any kind of alignment tool can be used to generate the MSA. The uploaded MSA can be provided in aligned FASTA, A3M, STOCKHOLM or Clustal format. We convert the respective MSA format into A3M format using the `reformat.pl` script from the HH-suite [8].

#### Lightweight 2D structure renderer

For visualization, we developed a matplotlib [32] compatible module for displaying the 3D ribbon diagram of a protein structure or complex. The ribbon can be colored by residue index (N to C terminus) or by a predicted confidence metric (such as pLDDT). For complexes, each protein can be colored by chain ID. Instead of using a 3D renderer, we instead use a 2D line plotting based technique. The lines that make up the ribbon are plotted in the order in which they appear along the z-axis. Furthermore, we add shade to the lines according to the z-axis. This creates the illusion of a 3D rendered graphic. The advantage over a 3D renderer is that the images are very lightweight, can be used in animations and saved as vector graphics for lossless inclusion in documents. As the 2D renderer is not interactive, we additionally included a 3D visualization using py3Dmol [33] in the ColabFold notebooks.

### Benchmarking ColabFold

We show with multiple datasets that ColabFold does not sacrifice accuracy for its much faster runtimes.

#### Benchmark with CASP14 targets

We compared AlphaFold-Colab and AlphaFold2 (commit b88f8da) against ColabFold using all CASP14 [2] targets. ColabFold-AlphaFold2 (commit `2b49880`) used UniRef30 (2021_03) [34] and the BFD/MGnify or ColabFoldDB. ColabFold-RoseTTAFold (commit `ae2b519`) was executed with papermill (github.com/nteract/papermill) using the PyRosetta protocol [28]. ColabFold-RoseTTAFold-BFD/MGnify and ColabFold-AlphaFold2-BFD/MGnify used the same MSAs. AlphaFold-Colab used the UniRef90 (2021_03), MGnify (2019_05) and the small BFD. AlphaFold2 used the `full_dbs` preset with and default databases downloaded with the `download_all_data.sh` script. The 65 targets contain 91 domains, among these are 20 FM-targets with 28 domains. We compared the predictions against the experimental structures using TMalign [35].

#### Measuring run-times for CASP14 benchmark

To provide more accurate run times we split MSA generation and model inference measurements. MSA generation times were repeated five times and averaged.

ColabFold was executed using `colabfold_batch`. The MMseqs2 server which computes MSAs for ColabFold has 2×14 core Intel E5-2680v4 CPUs and 768 GB RAM. Each generated MSA was processed by a single CPU-core. Runtimes were computed from server logs.

AlphaFold2 MSA generation runtimes were measured by running AlphaFold2 without models (providing an empty string to the `-- model_names` parameter) on the same 2×14 core Intel E5-2680v4 CPUs and 768 GB RAM system. The AlphaFold2 databases were stored on a software-RAID5 composed of six Samsung 970 EVO Plus 1TB NVMe drives. Runtimes for AlphaFold2 were taken from the `features` entry of the `timings.json` file. For a fair comparison, AlphaFold2 was modified to allow HMMer and HHblits to access one CPU core.

All ColabFold and AlphaFold2 model inference runtime measurements were done on systems with 2×16 core Intel Gold 6242 CPUs with 192 GB RAM and 4x Nvidia Quadro RTX5000 GPUs. Only one GPU was used in each run.

ColabFold-RoseTTAFold-BFD/MGnify and ColabFold-AlphaFold2-BFD/MGnify used the same MSAs, runtimes are shown only once.

AlphaFold-Colab was executed in the browser using a Google Colab Pro account. Times for homology search were taken from the notebook output cell “Search against genetic databases” cell. The JackHMMer search uses 8 threads.

#### Complex benchmark

We compare predictions of seventeen ClusPro [4, 12] targets to their native structures using DockQ [36]. We used `colabfold_batch` (commit 45ad0e9) with BFD/MGnify in residue-index manipulation- and AlphaFold-multimer mode to predict structures. We use MSA pairing as described in “MSA pairing for complex prediction” and also add unpaired sequences. Models are ranked by predicted interface pTMscore as returned by AlphaFold-multimer. The DockQ AlphaFold-multimer reference numbers were provided by Richard Evans.

#### Proteome benchmark

We predict the proteome of *M. jannaschii*. Of the 1787 proteins we exclude the 25 proteins longer than 1000 residues, leaving 1762 proteins of 268 aa average length. With the `colabfold_search` wrapper to MMseqs2 we search against the ColabFoldDB (“ColabFoldDB”) in 113 min on a system with an AMD EPYC 7402P 24-core CPU (no hyperthreading) and 512GB RAM. MMseqs2 had a maximum resident set size of 308 GB during the search. We then predict the structures on a single Nvidia Titan RTX with 24 GB RAM in 46 h using only MSAs (no templates). For each query we stop early if any recycle iteration reaches a pLDDT of at least 85. Early stopping results in a speed-up of 3.7× over default and 4.8× over always recompiling. AlphaFold2 (reduced_dbs) was ran with the reduced_dbs preset and no template information was used. We changed the AlphaFold2 source code to utilize all CPU cores during the homology search.

AlphaFold2 (reduced_dbs, v2.1.1), ColabFold (commit f5d0cec) default and ColabFold Stop ≥ 85 have an average pLDDT of 90.68, 90.22 and 89.33 respectively for 50 randomly sampled proteins. These are the same proteins that were used to extrapolate the run-time of AlphaFold2. Over all predictions, the pLDDTs for the *M. jannaschii* proteome downloaded from the AlphaFoldDB, ColabFold default and ColabFold Stop ≥ 85 are 89.75, 89.38 and 88.77, respectively.

### CODE AVAILABILITY

ColabFold is free open-source software (MIT) and available at github.com/sokrypton/ColabFold. A locally installable version is available at github.com/YoshitakaMo/ localcolabfold. The ColabFold development version shown in this manuscript is available at github.com/konstin/ColabFold. The ColabFold server components are free open-source software (GPLv3) and available at github.com/soedinglab/mmseqs2-app. MMseqs2 is free open-source software (GPLv3) and available at mmseqs.com.

### DATA AVAILABILITY

ColabFold databases are freely (CC-BY-SA 4.0) available at colabfold.mmseqs.com.

MSAs and structures produced during benchmarking: wwwuser.gwdg.de/~compbiol/colabfold/manuscript

Input databases used for building ColabFold databases: UniRef30: uniclust.mmseqs.com

BFD: bfd.mmseqs.com

MGnify: ftp.ebi.ac.uk/pub/databases/metagenomics/peptide_database/2019_05

PDB70: wwwuser.gwdg.de/~compbiol/data/hhsuite/databases/hhsuite_dbs

MetaEuk: wwwuser.gwdg.de/~compbiol/metaeuk/2019_11/MetaEuk_preds_Tara_vs_euk_profiles_uniqs.fas.gz

SMAG: www.genoscope.cns.fr/tara/localdata/data/SMAGs-v1/SMAGs_v1_concat.faa.tar.gz

TOPAZ: osf.io/gm564

MGV: portal.nersc.gov/MGV/MGV_v1.0_2021_07_08/mgv_ proteins.faa

GPD: ftp.ebi.ac.uk/pub/databases/metagenomics/genome_sets/gut_phage_database/GPD_proteome.faa

Further datasets used for benchmarking ColabFold: PFAM (Pfam-A.seed.gz & Pfam-A.full.gz): ftp.ebi.ac.uk/pub/databases/Pfam/releases/Pfam34.0

*M. jannaschii* proteome: http://uniprot.org/proteomes/UP000000805ftp.ebi.ac.uk/pub/databases/alphafold/v1/UP000000805_243232_METJA_v1.tar

## REFERENCES

[1] Jumper, J. et al. Nature 596, 583–589 (2021).

[2] Kryshtafovych, A. et al. Proteins 89, 1607–1617 (2021).

[3] Baek, M. et al. Science 373, 871–876 (2021).

[4] Evans, R. et al. bioRxiv 2021.10.04.463034 (2021).

[5] UniProt Consortium. Nucleic Acids Res. 47, D506–D515 (2019).

[6] Mitchell, A. L. et al. Nucleic Acids Res. 48, D570–D578 (2020).

[7] Eddy, S. R. PLoS Comput. Biol. 7, e1002195 (2011).

[8] Steinegger, M. et al. BMC Bioinform. 20, 473 (2019).

[9] Tunyasuvunakool, K. et al. Nature 596, 590–596 (2021).

[10] Steinegger, M. & Söding, J. Nat. Biotechnol. 35, 1026–1028 (2017).

[11] Mirdita, M. et al. Bioinformatics 35, 2856–2858 (2019).

[12] Kozakov, D. et al. Nat. Protoc. 12, 255–278 (2017).

[13] Levy Karin, E. et al. Microbiome 8, 48 (2020).

[14] Delmont, T. O. et al. bioRxiv 2020.10.15.341214 (2020).

[15] Alexander, H. et al. bioRxiv 2021.07.25.453713 (2021).

[16] Nayfach, S. et al. Nat. Microbiol. 6, 960–970 (2021).

[17] Camarillo-Guerrero, L. F. et al. Cell 184, 1098–1109.e9 (2021).

[18] Steinegger, M. & Söding, J. Nat. Commun. 9, 2542 (2018).

[19] Mistry, J. et al. Nucleic Acids Res. 49 (2021).

[20] Moriwaki, Y. AlphaFold2 can also predict heterocomplexes. all you have to do is input the two sequences you want to predict and connect them with a long linker. https://twitter.com/Ag_smith/status/1417063635000598528 (2021).

[21] Baek, M. Adding a big enough number for “residue_index” feature is enough to model hetero-complex using AlphaFold (green&cyan: crystal structure / magenta: predicted model w/ residue_index modification). https://twitter.com/minkbaek/status/1417538291709071362 (2021).

[22] Bryant, P. et al. bioRxiv 2021.09.15.460468 (2021).

[23] Mosalaganti, S. et al. bioRxiv 2021.10.26.465776 (2021).

## REFERENCES

[24] Kluyver, T. et al. Jupyter Notebooks - a publishing format for reproducible computational workflows. In Positioning and Power in Academic Publishing: Players, Agents and Agendas, 87–90 (IOS Press, 2016).

[25] Eastman, P. et al. PLoS Comput. Biol. 13, 1–17 (2017).

[26] Gal, Y. & Ghahramani, Z. arXiv:1506.02142 (2016).

[27] Krivov, G. G. et al. Proteins 77, 778–795 (2009).

[28] Chaudhury, S. et al. Bioinformatics 26, 689–691 (2010).

[29] Suzek, B. E. et al. Bioinformatics 31, 926–932 (2015).

[30] Federhen, S. Nucleic Acids Res. 40, D136–D143 (2012).

[31] Bradbury, J. et al. JAX: composable transformations of Python+NumPy programs (2018).

[32] Hunter, J. D. Comput. Sci. Eng. 9, 90–95 (2007).

[33] Rego, N. & Koes, D. Bioinformatics 31, 1322–1324 (2015).

[34] Mirdita, M. et al. Nucleic Acids Res. 45, D170–D176 (2017).

[35] Zhang, Y. & Skolnick, J. Nucleic Acids Res. 33, 2302–2309 (2005).

[36] Basu, S. & Wallner, B. PLoS One 11, e0161879 (2016).

