## Supplementary material for "ColabFold - Making protein folding accessible to all"

[1] Quantitative and Computational Biology, Max Planck Institute for Biophysical Chemistry, Göttingen, Germany. [2] School of Biological Sciences, Seoul National University, Seoul, South Korea. [3] Department of Biotechnology, Graduate School of Agricultural and Life Sciences, The University of Tokyo, Tokyo, Japan. [4] Collaborative Research Institute for Innovative Microbiology, The University of Tokyo, Tokyo, Japan. [5] Department of Biochemistry and Molecular Biology, Michigan State University, East Lansing, MI 48824, USA. [6] JHDSF Program, Harvard University, Cambridge, MA 02138, USA. [7] FAS Division of Science, Harvard University, Cambridge, MA 02138, USA. [8] Artificial Intelligence Institute, Seoul National University, Seoul, South Korea [*] These authors contributed equally and are ordered alphabetically.
**Contact:**

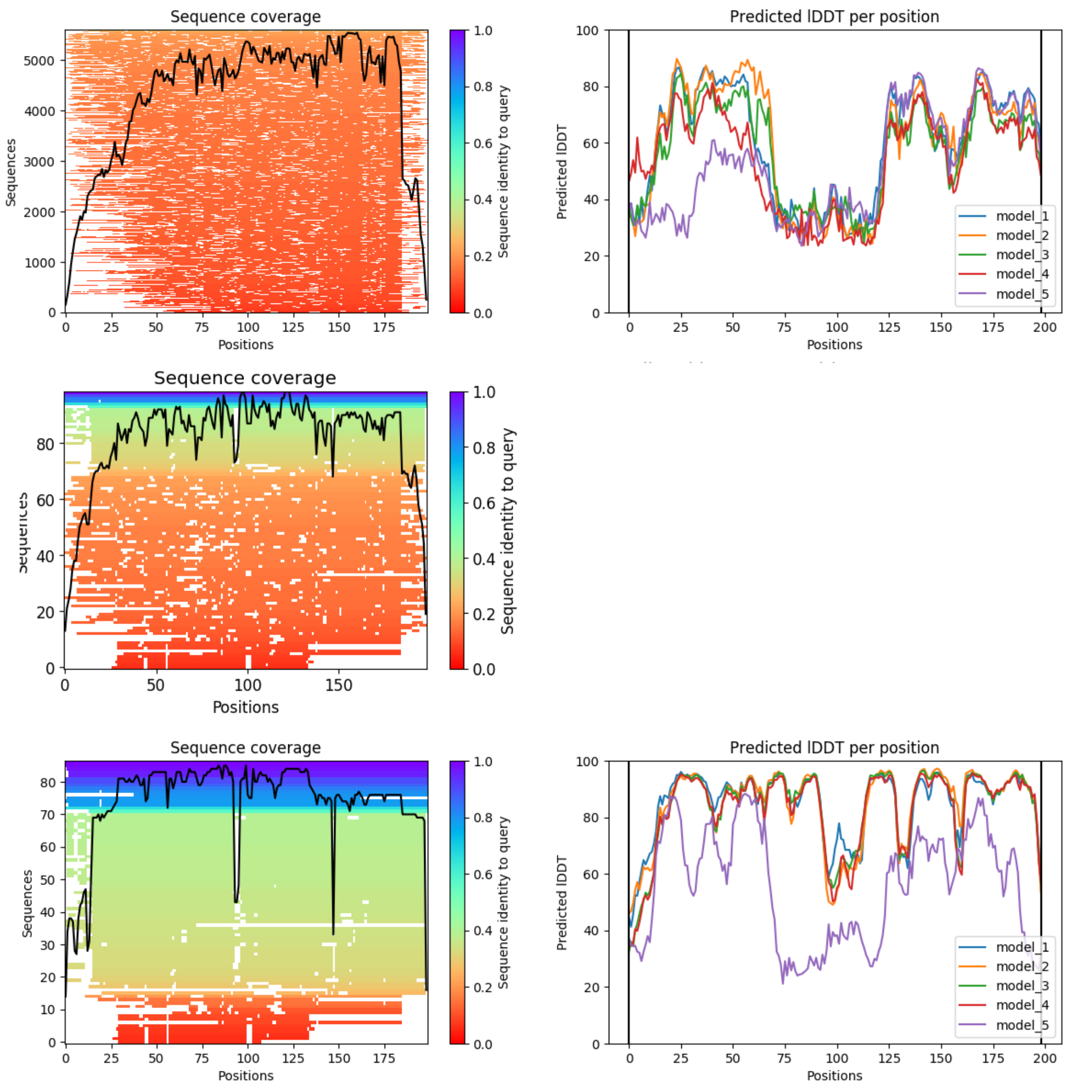


**Supplementary Figure** 1. **Anecdotal example of improved prediction through MSA filtering.** MSA coverage (left) and pLDDTs of predicted ColabFold models (right) for CASP14 target T1038 with two different filtering settings: Top: Single MSA filtering step with HHblits filtering algorithm and `--diff 3000` setting. Middle: Zoomed in view of first 100 sequences in top. Bottom: Three step MSA filtering as described in methods.

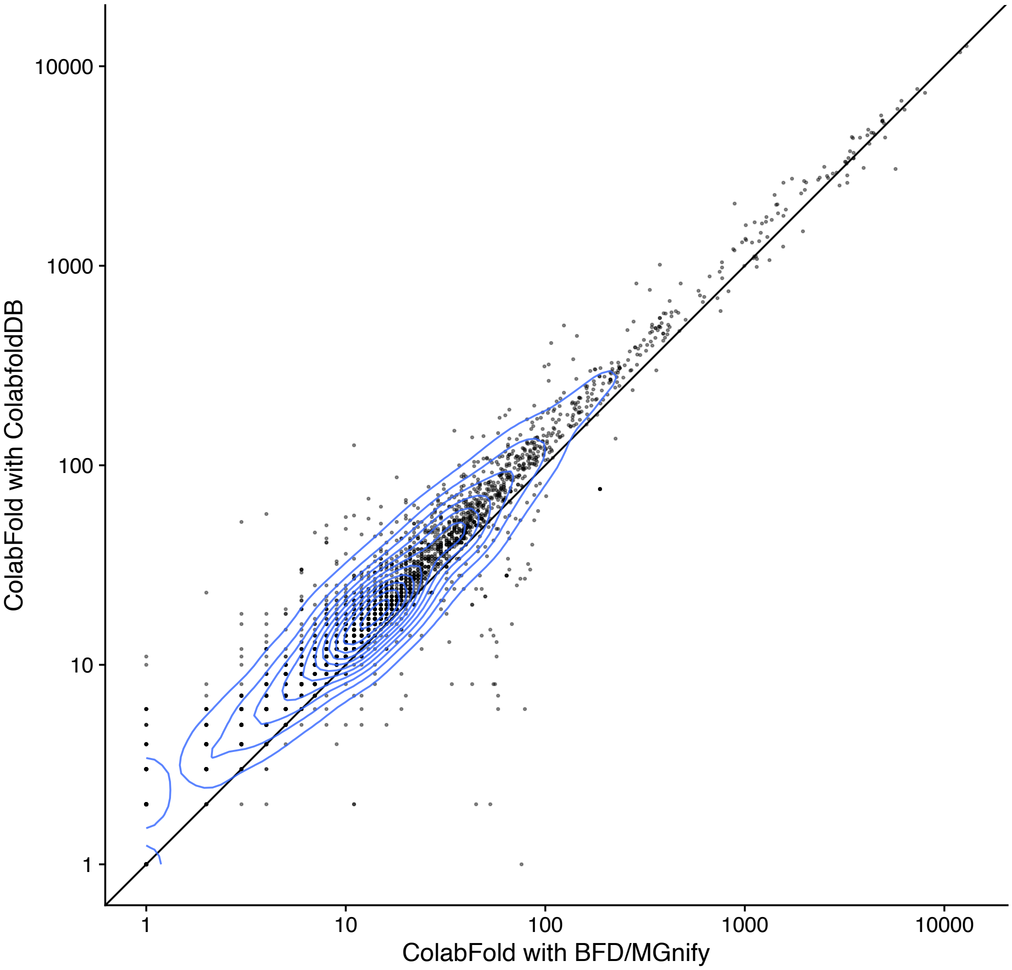


**Supplementary Figure** 2. **Comparison of Enrichment in PFAM** Comparison of homology search hits found from selected PFAM sequences against BFD/MGnify and ColabFoldDB. We select 2439 PFAM 34.0 entries that have less than 30 sequences in their `Pfam-A.full` entry. In each of these PFAM families we select from the `Pfam-A.seed` the longest sequence. We search this sequence with the ColabFold MMseqs2 workflow against the BFD/MGnify and ColabFoldDB. From the MSAs we cut the PFAM domains and note how many sequences cover the domain by at least 75%.

| | MSA Neff | | MSA #seq | |
|---|---|---|---|---|
| **Target** | **AlphaFold2** | **ColabFold** | **AlphaFold** | **ColabFold** |
| T1033-D1 | 1.8 | **2.2** | 5 | **6** |
| T1040-D1 | 4.7 | **5.5** | 38 | **74** |
| T1043-D1 | 1.7 | **5.6** | 5 | **57** |
| T1064-D1 | 2.1 | **2.3** | 9 | **13** |

**Supplementary Table** 1. **In-depth analysis of CASP14 targets that ColabFold predicted better than AlphaFold2.** The largest improvements were observed in four targets: (1) T1064-D1 is ORF8 from *SARS-CoV-2*, (2-4) T1033-D1, T1040-D1, T1043-D1 are single domains from a large RNA polymerase of the crAss-like phage. All of these target sequences are from the CASP14-FM category and lack homology even in large metagenomic databases like BFD or MGnify. We compared the MSAs by computing the Neff using `hhmake` from the HH-suite. Neff is an entropy measure for multiple sequences alignments, the larger the Neff the more diverse the MSA. Higher Neff values correlate with better AlphaFold2 predictions (see Jumper et al., Nature, 2021, **Fig. 5a**). The MMseqs2 search of ColabFold generates for all targets higher Neff values and therefore better predictions. In target T1033 a single additional sequence is enough to increase the TM-score from 0.348 (AlphaFold2) to 0.820 (ColabFold-AlphaFold2-BFD/MGnify). During CASP14 the AlphaFold team searched the RNA polymerase targets as a single sequence instead of separate domains, which resulted in much larger MSAs (see Jumper et al., Proteins, 2021, **Fig. 3**), while for our benchmark we searched each domain separately.

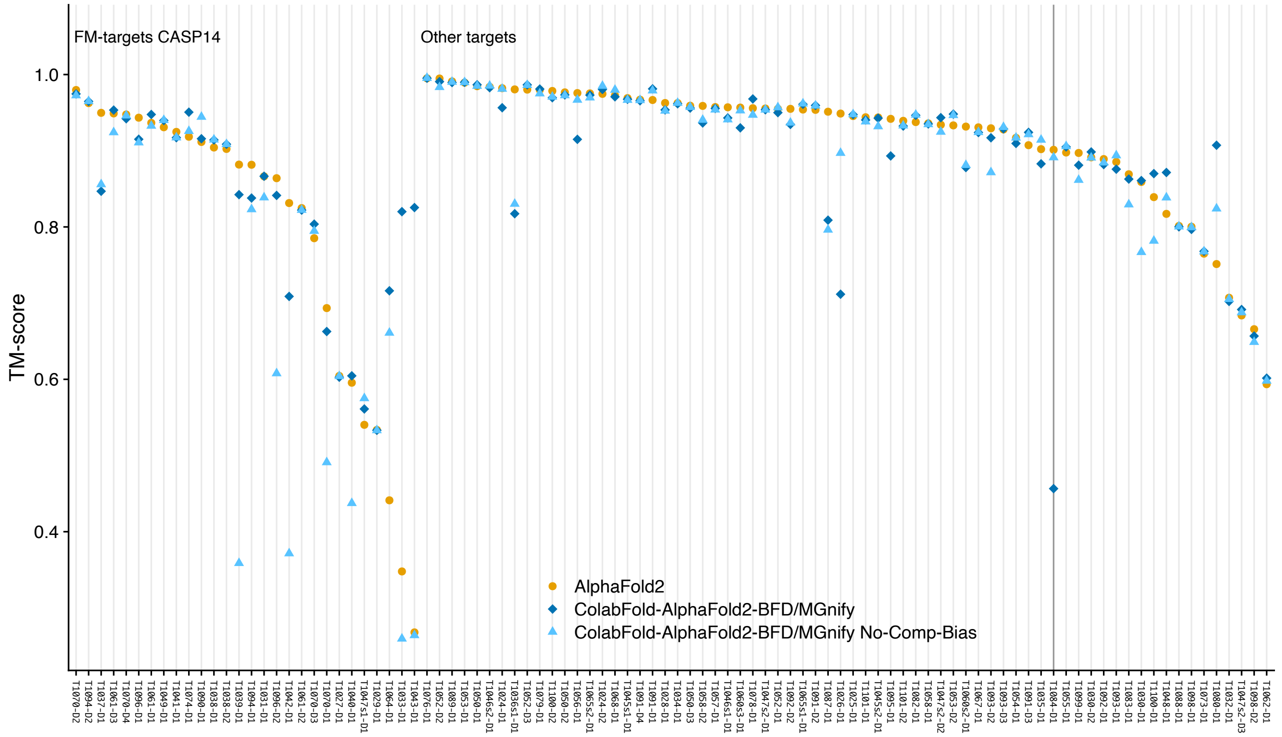


**Supplementary Figure 3. Disabling composition-bias and masking in MMseqs2 result in better accuracy for some CASP14 targets** We turned off two MMseqs2 mechanisms for false positive suppression (`--comp-bias-corr 0 --mask-profile 0`) and reran our CASP14 benchmark. Target T1084-D1 (highlighted) achieves now a TM-score of 0.891210 instead of 0.456540 in default search mode.

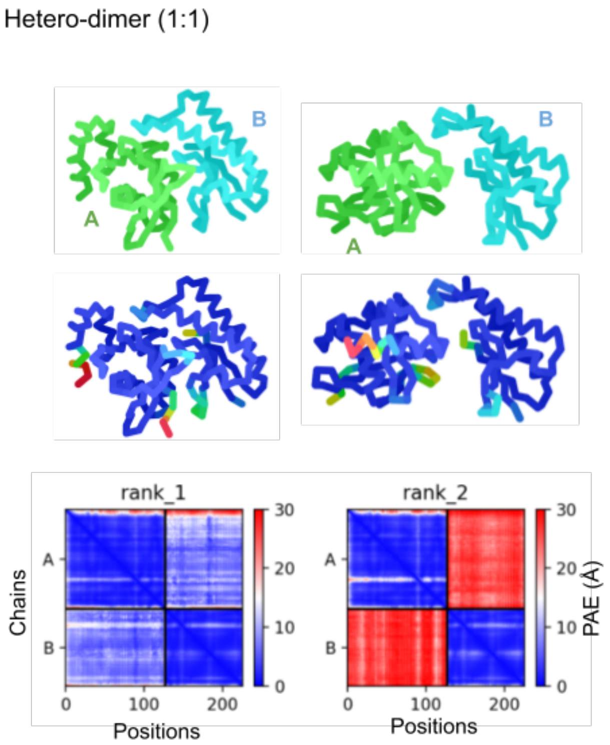


**Supplementary Figure** 4. **Hetero-dimer 1:1** Only one of the five models predicted for CASP14 target H1065 has a high agreement with its native structure during unpaired complex prediction. Although the pLDDT scores are nearly identical (shown in the middle with colored chains), the inter-PAE (bottom) is significantly lower (meaning more confident) for the correctly predicted complex (rank 1 vs rank 2). This demonstrates the utility of PAE (and the derived pTMscore) in ranking complexes.

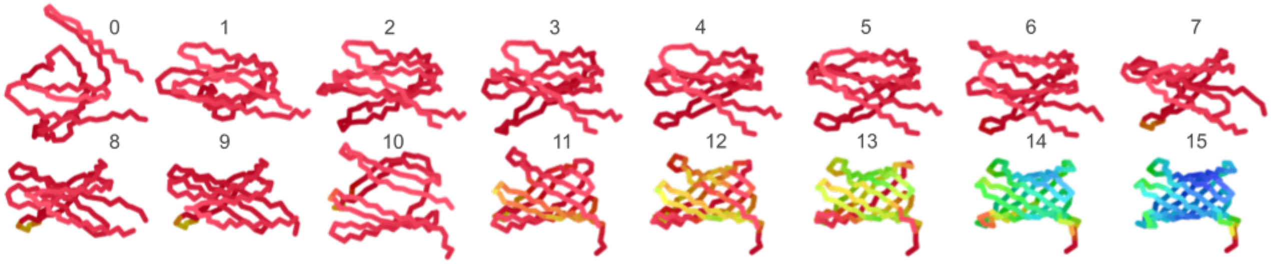


**Supplementary Figure** 5. **Example of additional recycle steps improving prediction** Occasionally, increasing the number of recycles can help find a well predicted structure. For this de-novo designed transmembrane protein (Vorobieva et al. Science, 371(6531), 2021), 15 recycle iterations were needed to produce structure with high pLDDT.

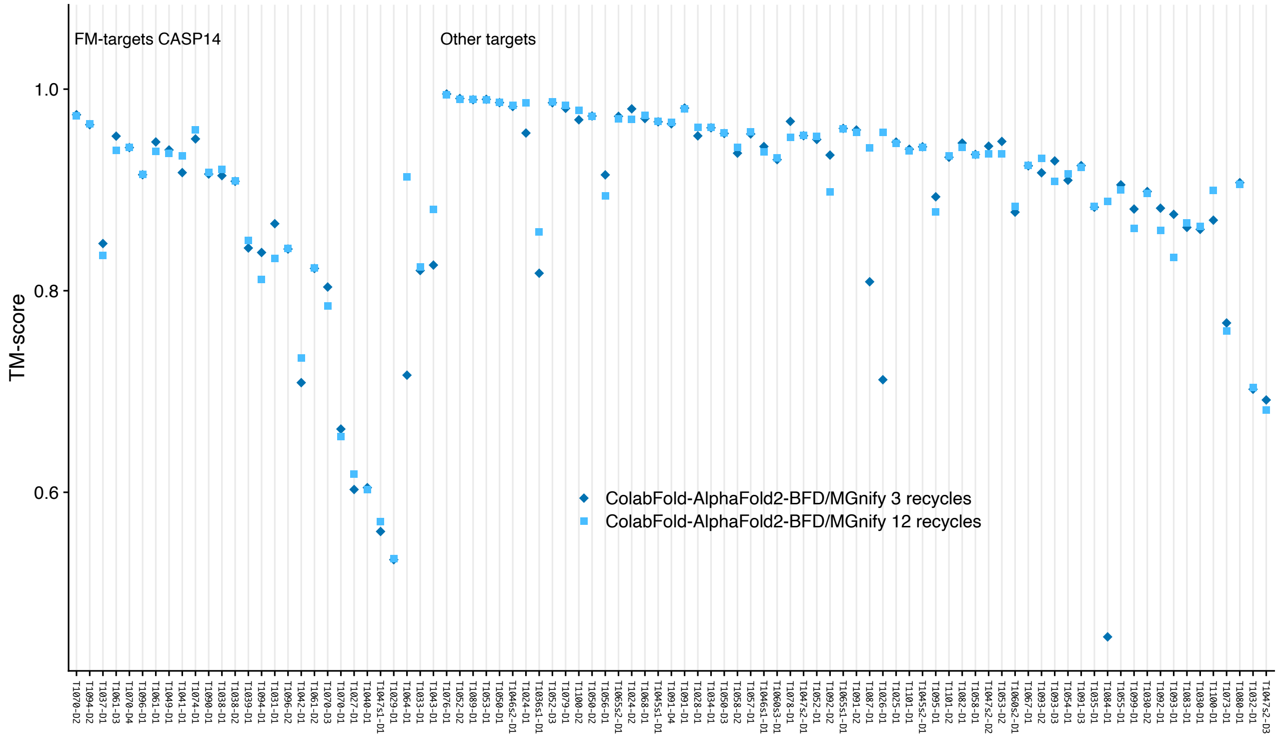


**Supplementary Figure** 6. **Increasing AlphaFold2 recycles from 3 to 12 results in better accuracy for some CASP14 targets** We increased the number of recycles executed by ColabFold-AlphaFold2 from 3 to 12 (`--num-recycle 12`) and reran our CASP14 benchmark.
